# Different subtypes of influenza viruses target different human proteins and pathways leading to different pathogenic phenotypes

**DOI:** 10.1101/541177

**Authors:** Yujie Wang, Ting Song, Kaiwu Li, Yuan Jin, Junjie Yue, Hongguang Ren, Long Liang

**Affiliations:** State Key Laboratory of Pathogen and Biosecurity, Beijing Institute of Biotechnology, Beijing 100071, China; Institute of Physical Science and Information Technology, Anhui University, Hefei, Anhui 230601, China

## Abstract

Different subtypes of Influenza A viruses (IAVs) cause different pathogenic phenotypes after infecting human bodies. Analysis of the interactions between viral proteins and the host proteins may provide insights into the pathogenic mechanisms of the virus. In this paper, we found that the same proteins (Nucleoprotein and Neuraminidase) of H1N1 and H5N1 have different impacts on the NF-κB activation. By further examining the virus-host protein-protein interactions, we found that both NP and NA proteins of the H1N1 and H5N1 viruses target on different host proteins. These results indicate that different subtypes of influenza viruses target different human proteins and pathways leading to different pathogenic phenotypes.

## Introduction

Influenza A virus (IAV) belongs to the Orthomyxoviridae family. Its genome consists of eight segmented, negative-strand RNA^[1–3]^. IAV are further typed into many different subtypes based on the antigenicity of hemagglutinin (HA) and the neuraminidase (NA)^[4]^, the proteins on the surface of the virus. Currently, 18 HA(H1-H18) and 11 NA (N1-N11) subtypes have been identified in IAVs^[5]^. Different subtypes of IAVs may have evolved various mechanisms to co-opt host processes and suppress host defenses, inducing different infectious phenotypic outcomes. For example, the highly transmissible seasonal H1N1 virus usually causes mild illness, while the highly pathogenic avian H5N1 virus often leads to severe cases when infecting humans. The pathogenic mechanism under this phenomenon needs more research efforts to reveal.

The innate immune system of human plays an important role in the hosts’ resistance to viruses. The NF-κB signaling pathway^[6–9]^ is a key component of the innate immune system during its antiviral process. In mammals, there are two major NF-κB signaling pathways: the canonical NF-κB pathway and the noncanonical NF-κB pathway^[10,11]^. Under the stimulation of pathogenic microbial infections or their induced inflammatory factors, NF-κB is normally activated by classical signaling pathways^[12]^. Diverse bacterial and viral pathogens target NF-κB signaling pathway to evade host immune defenses. Some viruses suppress NF-κB activation to dampen the host immune responses to maintain latency^[13]^. For example, human bocavirus (HBoV) proteins NS1 and NS1-70^[14]^, the mumps virus (MuV) small hydrophobic protein (SH)^[15]^ and Molluscum contagiosum virus (MCV) protein MC005^[16]^ inhibit TNF-α-mediated activation of NF-κB by reducing the phosphorylation of IKKβ, IκBα, and p65 as well as the translocation of p65 into the nucleus. Some pathogenic microorganisms activate NF-κB for viral gene expression, replication and spread. For example, the K15 protein of KSHV and the early protein Nef of most primate lentiviruses enhances NF-κB activation to initiate proviral transcription^[17,18]^. IAVs can also interfere with antiviral responses by regulating the NF-κB signaling pathway. Studies have shown that influenza virus proteins interact with NF-κB to promote viral replication. For example, the IAV NS1 protein specifically inhibits IKK-mediated NF-κB activation and production of the NF-κB induced antiviral genes by physically interacting with IKK through the C-terminal effector domain^[19]^.

In this paper, we used the luciferase assay and the Co-immunoprecipitation experiment to study how the NP and NA viral proteins of both HIN1 and H5N1 interact with NF-κB signaling pathway. We found that the H1N1 NP has little impact on the TNF-α-induced NF-κB transcriptional activation while the H5N1 NP inhibits the TNF-α-induced NF-κB transcriptional activation by interacting with IKKα. Furthermore, the H5N1 NP protein promotes the expression of IκBα proteins, meanwhile, suppresses the phosphorylation of IκBα, and restrains the translocation of p65 into the nucleus of infected 293T cells. We also found that the H5N1 NA actives the IL-1β-induced NF-κB transcriptional activation through TAB2, while H1N1 NA inhibits the IL-1β-induced NF-κB transcriptional activation by coimmunoprecipitating with IKKβ. All these results imply divergent pathogenic mechanisms of different subtypes of IAVs.

## Materials and Methods

### 1. Cell Culture and Transfection

293T (human embryonic kidney) and Hela cells were cultured in Dulbecco’s modified Eagle’s medium (DMEM, Gibco) supplemented with 10% heat-inactivated fetal bovine serum (FBS, Gibco), 2mM L-glutamine, 100 U/mL penicillin, and 100mg/mL streptomycin. Transient transfections were performed with Lipofectamine 2000 (Invitrogen) according to the manufacturer’s instructions.

### 2. Antibodies, Reagents

Anti-Flag (Sigma-Aldrich), anti-Myc (Santa Cruz), anti- *α* -tubulin (Sigma-Aldrich), and anti-HA (Origene) mouse monoclonal antibodies were used as primary antibodies for immunoblotting and immunoprecipitation. HRP-conjugated goat anti-mouse IgG (ZSGB-BIO) antibodies were used as secondary antibodies for immunoblotting.

### 3. Immunoprecipitation, and Immunoblotting

Immunoprecipitation was performed using IP buffer (1% Nonidet P-40, 50mM Tris-HCl [pH 7.5], 150mM NaCl, and Complete™ protease inhibitor cocktail-EDTA (Roche)). Whole cell extracts were prepared after transfection and incubated with indicated antibodies together with Protein A/G beads (Roche) overnight. Beads were then washed 4 times with IP buffer, and immunoprecipitates were eluted with SDS loading buffer (TransGen Biotech) and resolved in SDSPAGE gels. The proteins were transferred to PVDF membrane (Bio-Rad) and further incubated with the indicated antibodies. The antigen-antibody complexes were visualized by the Immubilon™ chemiluminescent detection kit (Millipore).

### 4. Dual-Luciferase Activity Assay

293T (human embryonic kidney) cells were seeded in 24-well plates (1 × 10^5^ per well). 293T (human embryonic kidney) cells were transfected with empty vector, 50 ng, 100 ng, 150 ng H5N1 NP or H1N1 NP expression plasmid, along with 125 ng pNF-κ B-luc, 25 ng pRL-TK utilizing transfection reagent (Invitrogen). After 24-hour transfection, luciferase activity was measured through the dual-luciferase assay system (Promega) in accordance with the manufacturer’s instructions.

### 5. Immunofluorescence analysis

HeLa cells cultured on coverslips were fixed in 4% paraformaldehyde for 10 min and washed with PBS for three times. After that, cells were fixed with 0.1% Triton X-100 on ice for 5min, washed in PBS and blocked in 1% BSA for 20 min. The coverslips were incubated with 1% BSA containing primary antibodies for 1 h followed by PBS wash for three times. The cells were further stained with Alexa Fluor^®^ conjugated secondary antibodies (Abcam). Images were acquired on an Olympus FV1000 fluorescence microscope.

### 6. Statistical Analysis

Analyses were done with the statistical software SAS/STAT. Data analysis over time was undertaken by repeated-measures analysis with SAS/STAT. Differences were considered statistically significant if the P value was <0.05.

## Results

### 1 H5N1 NP and H1N1 NP have different impacts on the TNF-α-induced NF-κB activation

The nucleoprotein of negative-strand RNA viruses forms a major component of the ribonucleoprotein complex that is responsible for viral transcription and replication, and NP proteins of IAV supports the progress of the polymerase during the elongation phase ^[20]^.

The dual-luciferase activity assay was performed to determine the impacts of the NP proteins from different subtypes of IAVs on the TNF-α-induced NF-κB activation. The results showed that the H5N1 NP clone significantly inhibited TNF-α stimulated NF-κB promoter activity, while the H1N1 NP had little impact on it (Figure 1a, Figure 1b).

**Figure 1:**
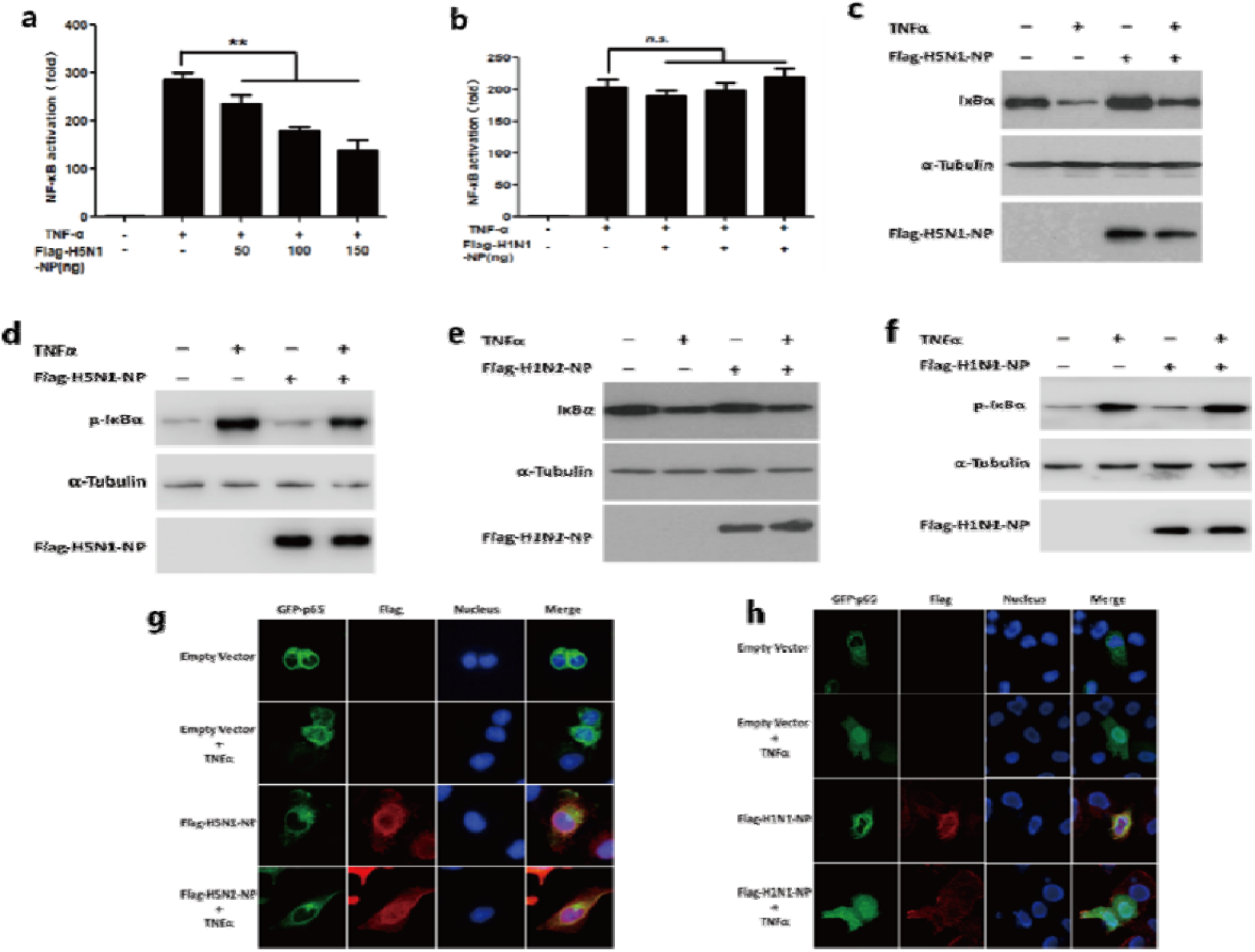
H5N1 NP and H1N1 NP have different impacts on TNF-α-induced NF-κB activation. 293T cells in 24-well plates were cotransfected with 125 ng pNF-κB-luc, 25 ng pRL-TK and indicated amount of H5N1 NP (a) and H1N1 NP (b) expression plasmid, or empty vector for 24 h. Cells were then mock-treated or treated with TNF-α (10 ng/ml) for 6 h. Reporter activity was determined by dual-luciferase reporter assays. The resultant ratios were normalized to the fold-change value by that of TNF-α -untreated cells cotransfected with empty vector, pNF-κB-luc and pRL-TK. Data shown represent three independent experiments, with each determination performed in duplicate (mean ± SD of fold change). Asterisks indicate significant differences between groups (**p < 0.05, Student’s t-test). 293T cells transfected with empty vector, H5N1 NP or H1N1 NP-expressing plasmid were stimulated with TNF-α (20 ng/ml) for indicated durations. Equal amounts of cell lysates were analyzed by immunoblotting with the anti-IκBα antibody or the anti-phospho-IκBα antibody(c-f). HeLa cells were transfected with H5N1 NP, H1N1 NP expression plasmid, or empty vector for 30 h. The cells were then mock-treated or treated with TNF-α (10 ng/ml) for 30 min. HeLa cells were subjected to immunofluorescence staining for detection of p65 subcellular localization by using rabbit anti–p65 and FITC-conjugated secondary Ab (green). H5N1 NP and H1N1 NP expression levels were detected using a mouse anti-Flag tag and Texas Red-conjugated secondary Ab (red). Nuclei were stained by Hoechst 33258 (blue) (g.h).

It is known that activated NF-κB signaling pathway can phosphorylate IκBα and cause subsequent degradation ^[21]^. To further verify the impact of the NP proteins on the NF-κB signaling pathway, we examined the phosphorylation and degradation of IκBα in the NF-κB signaling pathway by immunoblotting. When treated with TNF-α, the decreasing amount of IκBα in H5N1 NP-expressing cells was less than that in empty vector-transfected cells (Figure 1c). Furthermore, the level of phosphor-IκBα in the H5N1 NP-expressing cells was diminished (Figure 1d). The decreasing amount of IκBα in the H1N1 NP-expressing cells was similar to that in the empty vector-transfected cells, which were both treated with TNF-α (Figure 1e). The level of phosphor-IκBα in the H1N1 NP-expressing cells was similar to the empty vector-transfected cells (Figure 1f). These results demonstrated that H5N1 NP protein suppressed TNF-α -mediated IκBα phosphorylation and degradation while H1N1 NP protein had little impact on them.

The nuclear translocation of NF-κB is a prerequisite for the promotion of downstream genes^[22]^. Immunofluorescence also showed that almost all the p65 proteins were retained in the cytoplasm after treatment with TNF-α in the H5N1-NP-GFP-transfected cells (Figure 1g), while p65 transported from cytoplasm to nucleus in the H1N1-NP-GFP-transfected cells (Figure 1h). These findings indicated that H5N1 NP and H1N1 NP had different impacts on the TNF-α-induced NF-κB activation.

### 2 H5N1 NP inhibits the NF-κB signaling pathway by targeting IKKα

The results above indicated that H5N1 NP had an inhibitory effect on the NF-κB signaling pathway. Next, we intended to investigate which host protein the NP protein targeted on to get this effect. Firstly, we examined the effects of H5N1 NP protein on the luciferase activity mediated by overexpression of TNF-α signaling transducers along the NF-κB signaling pathway. H5N1 NP protein inhibited TRAF2, TAK1+TAB1, TAB2, IKKα-induced NF-κB activation in a dose-dependent manner (Figure 2a–2d) and had little effect on the IKKβ-mediated luciferase activity (Figure 2e). These results showed that the potential targets of H5N1 NP on the NF-κB signaling pathway may include the IKKα and the complex of TAK. Results of co-immunoprecipitation showed that H5N1 NP binds to IKKα, indicating that there were protein-protein interactions between H5N1 NP and the TAK1 and IKKα in cells (Figure 2f).

**Figure 2:**
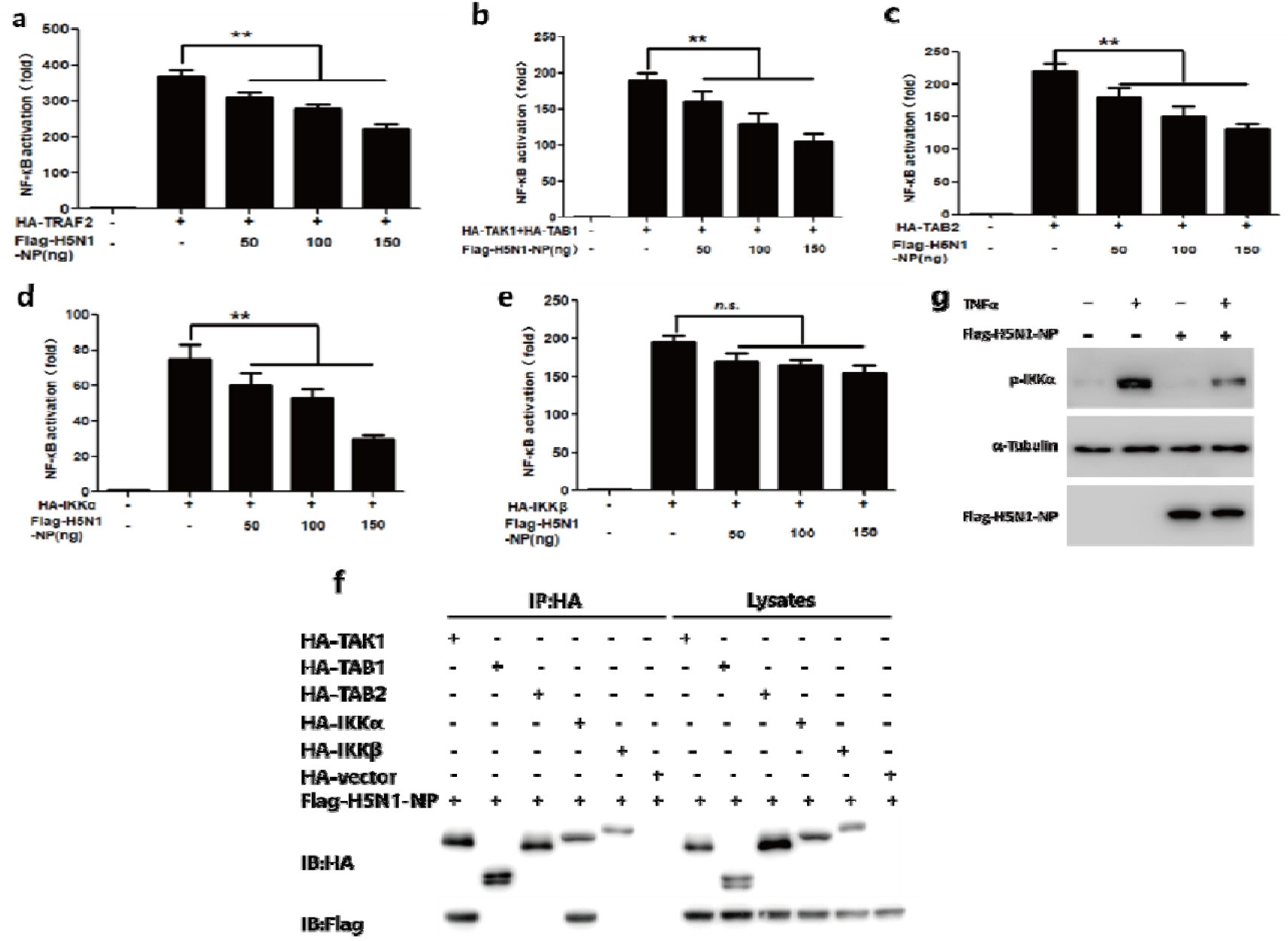
H5N1 NP inhibits the NF-κB signaling pathway by targeting IKKα. 293T cells in 24-well plates were cotransfected with 125 ng of pNF-κB-luc, 25 ng of pRL-TK and either HA-TRAF2 (a), HA-TAK1+HA-TAB1(b), HA-TAB2(c), HA-IKKα (d), HA-IKKβ (e) together with the indicated amounts of H5N1 NP expression plasmids. Total amounts of transfected DNA were kept equal by adding empty vector. Reporter activity was determined 30 h post-transfection by the dual-luciferase reporter assays. The resultant ratios were normalized to the foldchange value by that of cells cotransfected with empty vectors, pNF-κB-luc and pRL-TK. Data represent at least 3 independent experiments, with each determination performed in duplicate (mean ± SD of fold-change). Asterisks indicate significant differences between groups (** p < 0.05, Student’s t-test). 293T cells were transfected with Flag-H5N1 NP or either HA-TAK1 or HA-TAB1 or HA-TAB2 or HA-IKKα or HA-IKKβ or HA-vector expression plasmids for 30 h. All cells were then treated with TNF-α (10 ng/ml) for 30 min. The cells were lysed and subjected to immunoprecipitation (IP) using the mouse anti-HA tag. IP products and 5% input samples were analyzed by immunoblotting (f). 293T cells transfected with empty vector, H5N1 NP-expressing plasmid were stimulated with TNF-α (20 ng/ml) for indicated durations. Equal amounts of cell lysates were analyzed by immunoblotting with the anti-phospho-IKKα antibody (g).

The activation of IKKα by phosphorylation is required to the phosphorylation of IκB^[23]^. We therefore tested whether H5N1 NP played a role in inhibiting the phosphorylation of the IKKα. The results showed that H5N1 NP protein may have blocked the TNF-α-mediated IKKα phosphorylation(Figure 2g).

### 3 H5N1 NA and H1N1 NA have different impacts on the IL-1β-induced NF-κB activation

Neuraminidase(NA) is an integral membrane glycoprotein and a second major surface antigen of the IAV. During the infection process of IAV, NA cleaves terminal sialic acid from glycoproteins or glycolipids to free virus particles from host cell receptors and facilitate virus spread^[6,24]^.

To study the different impacts of the NA proteins of H5N1 and H1N1 on the NF-κB signaling pathways, reporter plasmid pNF-κB-luc and internal control plasmid pRL-TK, together with pH5N1 NA and pH1N1 NA or empty vector, were cotransfected into 293T cells. At 24h post-transfection, cells were mock-treated or treated with human IL-1β for 6h. These results showed that the H5N1 NA significantly promoted IL-1β-stimulated NF-κB promoter activity (Figure 3a), while the H1N1 NA had an inhibitory impact on it (Figure 3b).

**Figure 3:**
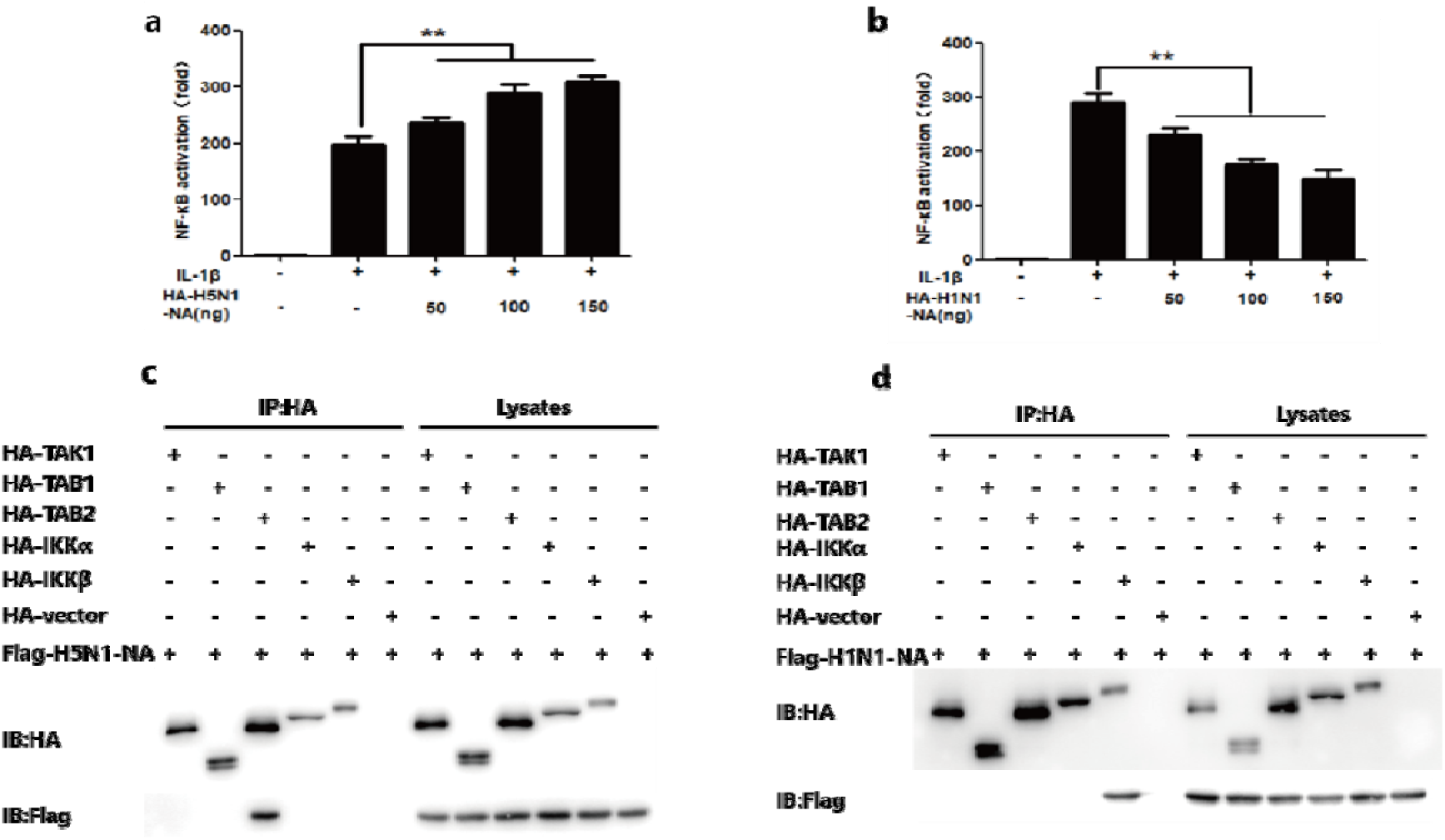
H5N1 NA and H1N1 NA have different impacts on IL-1β-induced NF-κB activation. 293T cells in 24-well plates were cotransfected with 125 ng pNF-κB-luc, 25 ng pRL-TK and indicated amount of H5N1 NA (a) and H1N1 NA(b) expression plasmid, or empty vector for 24 h. Cells were then mock-treated or treated with TNF-α (10 ng/ml) for 6 h. Reporter activity was determined by dual-luciferase reporter assays. The resultant ratios were normalized to the fold-change value by that of TNF-α -untreated cells cotransfected with empty vector, pNF-κB-luc and pRL-TK. Data shown represent three independent experiments, with each determination performed in duplicate (mean ± SD of fold change). Asterisks indicate significant differences between groups (**p < 0.05, Student’s t-test). 293T cells were transfected with Flag-H5N1 NA(c) or Flag-H1N1 NA(d) or either HA-TAK1 or HA-TAB1 or HA-TAB2 or HA-IKKα or HA-IKKβ or HA-vector expression plasmids for 30 h. All cells were then treated with TNF-α (10 ng/ml) for 30 min. The cells were lysed and subjected to immunoprecipitation (IP) using the mouse anti-HA tag. IP products and 5% input samples were analyzed by immunoblotting.

We inferred that the potential targets of the H5N1 NA on the NF-κB signaling pathway was the complex of TAK (Figure S1), and the potential targets of the H1N1 NA were TAB2 and IKKβ (Figure S2). Similarly, by implementing Co-Immunoprecipitation essays, we found that H5N1 NA interacted with TAB2 of the NF-κB signaling pathway (Figure 3c), while H1N1 NA interacted with IKKβ in the NF-κB signaling pathway (Figure 3d). Based on these results, we conclude that H5N1 NA and H1N1 NA had different impacts on the TNF-α-induced NF-κB activation.

## Discussion

NF-κB signaling pathway is widely found in almost all animal cells and plays a key role in the innate immune defense of host against pathogenic microorganisms invaded^[25]^. Once being invaded by viruses, the host manipulates the NF-κB signaling pathway to clear them. To survive, microbial pathogens have evolved strategies to modulate the NF-κB signaling pathway, which is an important pathogenic mechanism to evade innate immunity and promote replication.

In the process of influenza virus infecting the host, viral proteins or other pathogenic factors will interact with host proteins and signaling pathways. Previous research found that the IAV NS1 protein specifically inhibits IKK-mediated NF-κB activation and production of the NF-κB induced antiviral genes^[19]^. One report showed that PB1-F2 binds to IKKβ and impairs DNA-binding of NF-κB ^[26]^. In contrast, another study reports on an NF-κB intensifying activity of PB1-F2^[27]^. In our study, we focused on the regulation of NF-κB signaling pathway by NP and NA protein.

Firstly, we found the NP protein of H5N1 virus inhibited the transcriptional activation of the TNF-α-induced NF-κB signaling pathway (A/goose/Jilin/hb/2003 (H5N1), Figure 1a)^[28]^. In the absence of stimuli, NF-κB dimers are retained in the cytosol through association with an inhibitor of κB activity, termed IκB ^[29]^. Stimulus-induced degradation of IκB molecules leads to nuclear accumulation of p65, which is the basis of NF-κB activation. In our study, H5N1 NP protein abolished IkBa phosphorylation and degradation of IκBα, which making NF-κB dimers immobilized in the cytoplasm. The results of immunofluorescence experiments in Figure 1 verified that nuclear factors were still in the cytoplasm. Our study reveals a potential mechanism by which H5N1 virus evades human innate immune responses.

However, in our experiment with HIN1 NP as the experimental object, we found that H1N1 NP hardly affected the NF-κB signaling pathway (A/Beijing/501/2009 (H1N1), Figure 1b). We did not observe any significant change in the degradation and phosphorylation of IκBα protein. Nuclear translocation of p65 was also not affected by H1N1 NP. We speculate that the different effects between the two viral NP proteins and the NF-κB signaling pathway lead to different pathogenicity to some extent, but a more detailed mechanism needs further investigation.

NA is one of the major antigenic targets of the humoral immune response to IAV and the target of the antiviral drugs oseltamivir and zanamivir ^[30]^. Our studies on the neuraminidase protein showed that the NA protein of H5N1 virus promotes the transcriptional activation of IL-1β-induced NF-κB, while the NA protein of H1N1 influenza virus inhibits the corresponding activation.

The comparison of the above two groups of results indicates that the interaction between influenza virus proteins and host signaling pathway proteins has subtype-specific characteristics, which can respond to the specific pathogenic mechanism of influenza virus of different subtypes. The viral proteins employed to target at these conserved pathways are different for different subtypes of IAVs, suggesting more subtype-specific pathogenic mechanisms need to be revealed in the future research work.

## Conclusion

Our study shows that different proteins of varied influenza viruses will produce distinct interaction effects with host NF-κB signaling pathway. The interaction between influenza viral proteins and host proteins is subtype-specific. This specificity may due to the specifically pathogenic mechanisms of different subtypes of influenza viruses. Our discovery may suggest a new way of thinking about flu treatment and provide potential avenues to study pathogenesis of avian influenza.

## Acknowledgments

This work was supported by the National Natural Science Foundation of China (31800136), Beijing Nova Program (Z171100001117120) and the Independent Research Project from State Key Laboratory of Pathogen and Biosecurity (SKLPBS1807).

## Author contributions

Long Liang, Hongguang Ren and Junjie Yue conceived and supervised the study. Ting Song and Long Liang designed the experiments. Yujie Wang, Ting Song and Kaiwu Li performed the experiments. Hongguang Ren and Yuan Jin analyzed the data. Hongguang Ren and Yujie Wang wrote the manuscript. Yujie Wang and Ting Song contributed equally to this work.

## Supplementary Materials

Figure S1 and Figure S2 are the supplementary material, which include the the potential targets of the H5N1 NA and H1N1 NA in NF-κB signaling pathway. (*Supplementary Materials*)

