## supplementary material for "Different subtypes of influenza viruses target different human proteins and pathways leading to different pathogenic phenotypes"

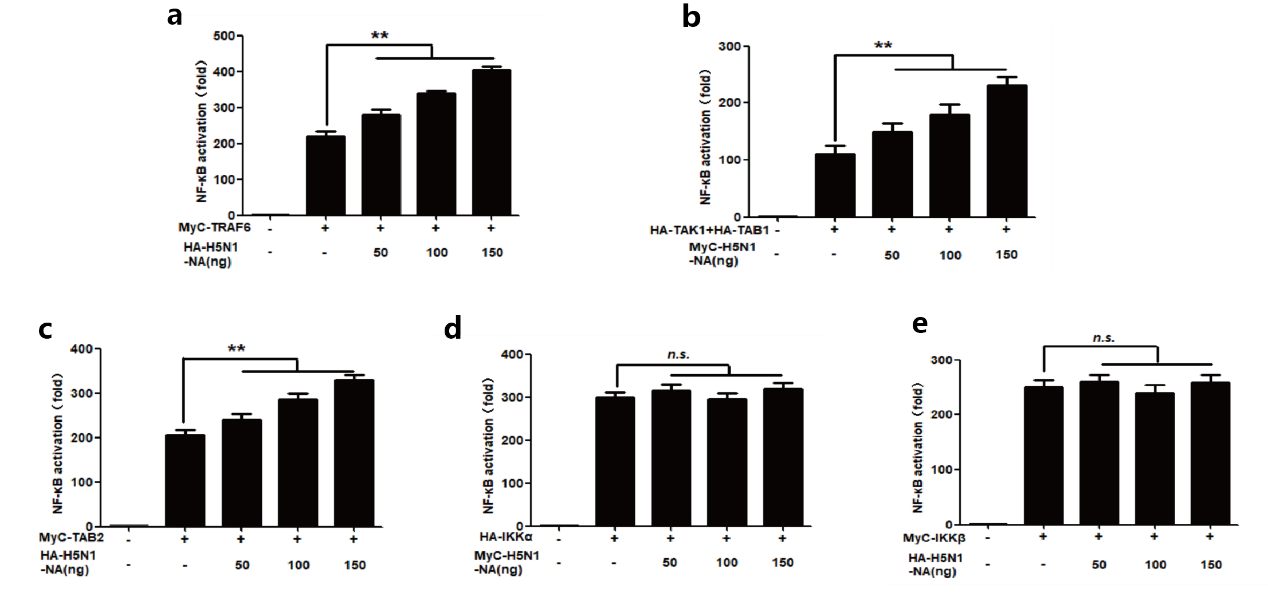

Figure S1: The potential target of H5N1 NA on the NF-κB signaling pathway is the complex of TAK. 293T cells in 24-well plates were cotransfected with 125 ng of pNF-κB-luc, 25 ng of pRL-TK and either HA-TRAF2 (a), HA-TAK1+HA-TAB1(b), HA-TAB2(c), HA-IKKα (d), HA-IKKβ (e) together with the indicated amounts of H5N1 NA expression plasmids. Total amounts of transfected DNA were kept equal by adding empty vector. Reporter activity was determined 30 h post-transfection by the dual-luciferase reporter assays. The resultant ratios were normalized to the foldchange value by that of cells cotransfected with empty vectors, pNF-κB-luc and pRL-TK. Data represent at least 3 independent experiments, with each determination performed in duplicate (mean ± SD of fold-change). Asterisks indicate significant differences between groups (** p < 0.05, Student’s t-test).

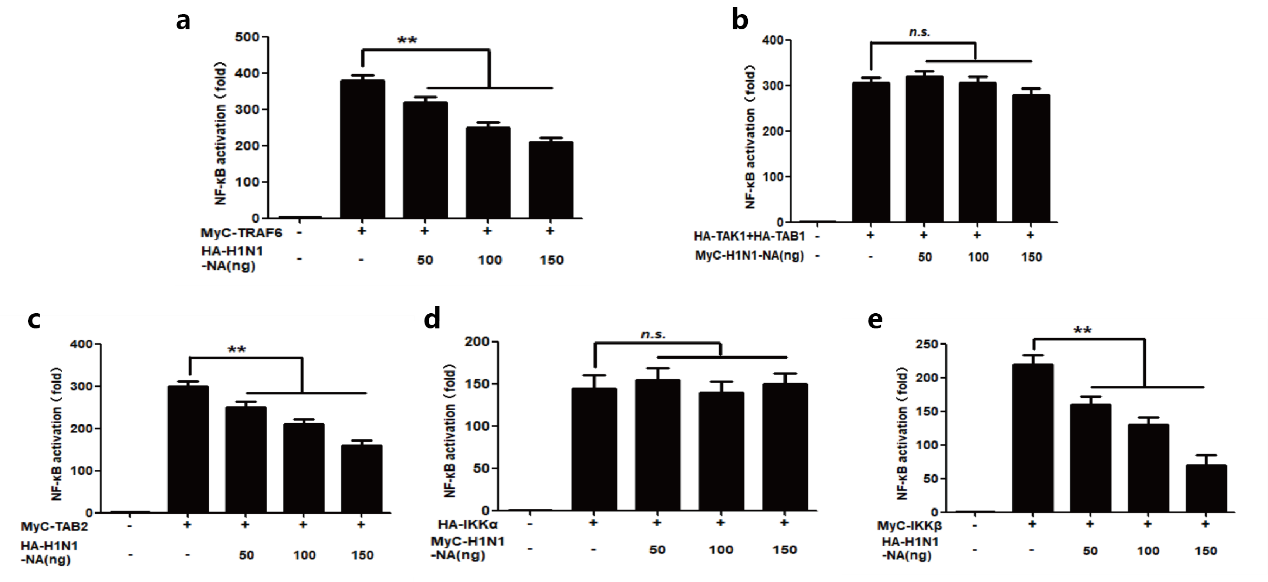

Figure S2: The potential targets of H1N1 NA on the NF-κB signaling pathway are TAB2 and IKKβ.

293T cells in 24-well plates were cotransfected with 125 ng of pNF-κB-luc, 25 ng of pRL-TK and either HA-TRAF2 (a), HA-TAK1+HA-TAB1(b), HA-TAB2(c), HA-IKKα (d), HA-IKKβ (e) together with the indicated amounts of H1N1 NA expression plasmids. Total amounts of transfected DNA were kept equal by adding empty vector. Reporter activity was determined 30 h post-transfection by the dual-luciferase reporter assays. The resultant ratios were normalized to the foldchange value by that of cells cotransfected with empty vectors, pNF-κ B-luc and pRL-TK. Data represent at least 3 independent experiments, with each determination performed in duplicate (mean ± SD of fold-change). Asterisks indicate significant differences between groups (** p < 0.05, Student’s t-test).
